# Co-occurrence of organic and inorganic N sources influences asparagine uptake and internal amino acid profiles in white clover

**DOI:** 10.1101/577114

**Authors:** Weronika Czaban, Jim Rasmussen

## Abstract

Direct plant uptake of organic nitrogen (N) is important for plant N nutrition, but we lack knowledge of how the concentration of external N forms (organic and inorganic) -influence organic N uptake and plant N status. We investigated the uptake of the amino acid asparagine (Asn) in white clover in the presence of different nitrate (NO_3_^-^), Asn, and total N concentrations. White clover seedlings were for one week exposed to combinations of NO_3_^-^ (3-30 µmol N kg^-1^ sand DW) and Asn (3-30 µmol N kg^-1^ sand DW), where after the Asn uptake rate was determined by addition of ^13^C_4_-Asn. Shoot and root Asn content and amino acid profiles were also analyzed. Increasing external NO_3_^-^ and total N concentrations decreased ^13^C_4_-Asn uptake rates and internal clover Asn content. Furthermore, total N affected clover amino acid profiles from non-essential amino acids at low N doses to the dominance of essential amino acids at increasing N doses. Asn uptake rate in white clover is reduced by increasing inorganic N, but not by increasing organic N concentrations. Furthermore, plant amino acid profiles are likely to be a more sensitive indicator of N supply and descriptor of the N status.

## Introduction

Legumes (*Leguminosae*) are of great significance to human, food, and animal feed due to their high nitrogen content and are mainly grown as grain, and forage species (1). They are rich in amino acids that are assimilated or derived from N accessed in three different processes: (1) N_2_-fixation, (2) inorganic N uptake (NO_3_^-^, NH_4_^+^), (3) organic N uptake (amino acids) (2, 3). N_2_-fixation and inorganic N uptake have been widely studied in legumes (4-6). In terms of energy cost, amino acid assimilation through these two pathways is the most-energy demanding. While direct uptake of organically bound N, where N is already in reduced form, costs less energy (7). Moreover, carbon cost of assimilating organic N into proteins is lower than that of inorganic N, mainly because of its carbon content. This carbon bonus makes it more beneficial for plants to take up organic than inorganic N (8). We recently reported white clover (*Trifolium repens*, cv. Rivendel) uptake of Asn in a sterile hydroponic solution (3) and in soil conditions at field relevant concentrations (9). Other legumes such as alfalfa (*Medicago sativa), alsike-clover (Trifolium hybridum L., cv. Stena), and red clover (T. pratense L., cv.Betty) have also been reported to take up organic N in laboratory or soil studies* (10, 11). Although the literature underlines the potential of amino acid absorption by legumes, the importance of amino acid uptake is unclear. This uncertainty is due to a lack of knowledge on how the amino acid uptake is influenced by the presence of other N forms that occur in soil simultaneously and can be acquired by N_2_-fixing legumes. To advance our understanding on how much amino acids contribute to the legume N budget, plant growth should be compared in soils that differ in N quality and quantity.

Information on the co-occurrence of different N forms and their influence on root absorption of amino acids mainly comes from studies with non-legume crop and tree species. One of the common findings is that uptake of amino acids is increased, while the absorption of inorganic N is reduced in mixtures of different N sources. Perennial ryegrass (*Lollium perenne*) exposed to a single and equimolar mixture of N sources (2 m*M* total N) doubled the uptake of glycine when supplied with NO_3_^-^ and NH_4_^+^ compared to when supplied alone (12). Spring wheat (*Triticum aestivum* L. cv. Amaretto) downregulated the assimilation of NO_3_^-^ when exposed to a mixture of glycine (1 m*M*) and NO_3_^-^ (1 m*M*), while the acquisition of the amino acid was unaffected by the NO_3_^-^ (13). Pre-incubation of young spruce (*Picea abies* L.) and beech (*Fagus sylvatica* L.) with amino acids (10 m*M*) reduced the NO_3_^-^ uptake when exposed to NO_3_^-^ and NH_4_^+^ (300-600 µ*M*), whereas the root amino acid content increased (14). Similar results were obtained for non-mycorrhizal beech (*Fagus sylvatica* L.), which was fed with NO_3_NH_4_ with and without amino acids: uptake of organic N was significantly higher than inorganic N (15). On the contrary, other studies on temperate and tropical forests point to a preferential uptake of inorganic rather than organic N from mixtures different N sources (16, 17). Whether such interactions occur in legumes is unknown. However, a significance of amino acid uptake in the presence of inorganic N has been documented for white clover (*Trifolium repens*). Based on the detection of L-asparagine-^13^C_4_,^15^N_2_, we reported uptake rates of 0.4 and 0.04 µmol g^-1^ root DW h^-1^ in a sterile hydroponic solution in both the presence and absence of NH_4_NO_3_ (3). Clover plants were also shown to compete for amino acids under soil conditions with uptake rates between 0.05 and 0.15 µmol g^-1^ root DW h^-1^ (9). Hence, amino acids can constitute a significant portion of N that is acquired by legumes, but we lack knowledge on the uptake of organic N in the presence of different N mixtures.

N uptake is thought to be strongly regulated by the N demand of the plant, specifically by the pool of amino compounds circulating between shoot and roots (7). Correspondingly, changes in amino acid composition can be affected by the N supply (18). This was observed by Cambui et al. (2011) who reported that in *Arabidopsis thaliana* grown on a mixture with NO_3_ and glutamine, a greater fraction of root N was derived from organic than inorganic N. Some studies also reported that plants supplied with organic N show different root morphology and higher root:shoot ratio than those supplied with inorganic N (8). A relation between amino acid composition and N supply was also found in legumes. We demonstrated for *Trifolium repens* that N supplementation (ON vs. ON + IN) affected the abundance of amino acids in the shoots, whereas in the roots, only the concentration (10 μ*M* vs. 1 m*M*) influenced the amino acid profile (3). This study indicated that root metabolism is more sensitive to nutrient quantity than quality. Thus understanding how the co-occurrence of different N forms in soil solution affects legume root and shoot performance would be fundamental not only from a pure scientific perspective, but also for the productivity and quality of forage legumes in agroecosystems. A complex interaction between the organic and inorganic N suggests that the co-occurrence of different N forms, rather than the presence of one, affects plant N uptake. However, so far this hypothesis has not been tested for legumes. Moreover, despite the extensive collection of data, studies mostly determined the uptake of amino acids from a one defined N mixture (12, 13, 17), while in soil solution inorganic and organic N occur in mixtures at various ratios and concentrations. Therefore, in this study the objective was to determine how different external N combinations influence and regulate amino acid uptake in a N_2_-fixing legume, white clover (*Trifolium repens*, cv. Rivendel). To systematically address some of the possible occurrences of organic and inorganic N in soil solution, we used an experimental space of N treatments. Three mechanisms are proposed for the external N regulation of amino acid uptake (Fig 1a):

1. a single N source: ON or IN independent of the presence of the other one,
2. the IN/ON ratio,
3. neither IN or ON, but the total N.

**Fig 1.**
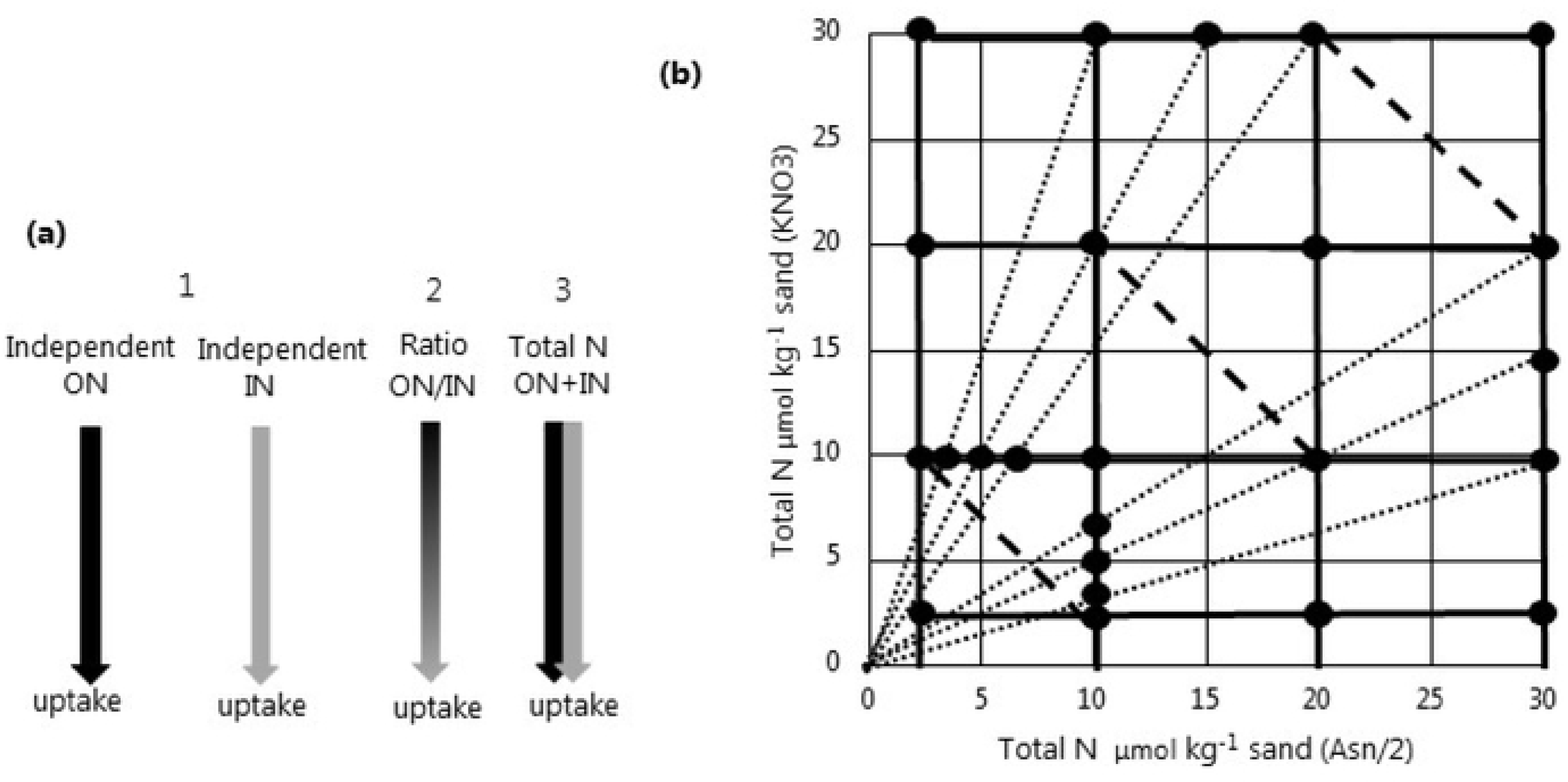
Experimental design. (a) Hypothetical models to explain regulation of Asn uptake, model 1 represents a single N source: ON or IN independent of the presence of the other one, model 2 represents ON/IN ratio, and model 3 represents the total N. (b) Matrix of 24 treatments to investigate possible regulation effects on Asn uptake according to model 1 (solid line), 2 (dotted line), and 3 (dashed line), each point on the graph represents one treatment (n = 4), the x-axis indicates the concentration of organic N (Asn), the y-axis indicates the concentration of inorganic N (KNO_3_)

White clover was chosen due to importance of amino acid uptake in this plant in both hydroponic and soil conditions (3, 9). The uptake of amino acid was evaluated based on acquisition of Asn, which is known to be taken up by clover and to be the most abundant amino acid in clover root extracts (19, 20).

## Materials and methods

### Plant and Rhizobium material

Seeds of white clover (*Trifolium repens* L., cv. Rivendel) were surface sterilized with sodium hypochlorite, rinsed with water, and germinated in filter centrifugal tubes (25 ml, Thermo Scientific) containing 38 g of inert Ottawa sand and placed in the climate chamber (day/night temperature of 18/8 °C; 16 h photoperiod of 70 μmol m^-2^ s^-1^). All seedlings received basic nutrient solution (2 ml) modified based on Varin, Cliquet (19) (mM): (mM): 0.18 CaCO_3_, 0.4 KH_2_PO_4_, 0.15 K_2_HPO_4_, 3 CaCl_2_, 0.375 MgSO_4_, 0.2 EDTA 2NaFe(3H_2_O), 0.014 H_3_BO_4_, 0.003 ZnSO_4_ x 7H_2_O, 0.0007 CuSO_4_, 0.117 Na_2_MoO_4_, 0.0001 CoCl_2_, 0.005 MnSO_4_, containing KNO_3_ and Asn as an inorganic and organic N source, respectively. Seedlings were assigned to testing and control groups: (1) 96 seedlings were assigned to testing group, which received basic nutrient solution containing ^15^N-KNO_3_ (3 µmol N kg^-1^ sand) and ^15^N-Asn (3 µmol N kg^-1^ sand). KNO_3_ and Asn were added as ^15^N-labeled (both at 5 at%) for later assessment of N_2_-fixation by the isotope dilution method (21). Five, seven, and nice days after sowing (DAS), those seedlings were inoculated with *Rhizobium leguminosarum* bv. trifolii TA1, (2)four uninoculated seedlings were assigned as control for the isotope dilution method and received basic nutrient solution containing ^15^N-KNO_3_, 5 at% (30 µmol N kg^-1^ sand) and ^15^N-Asn, 5 at% (30 µmol N kg^-1^ sand), (3) four uninoculated seedlings were assigned as control for the natural abundance of ^15^N and received basic nutrient solution containing ^14^N-KNO_3_ (30 µmol N kg^-1^ sand) and ^14^N-Asn, (30 µmol N kg^-1^ sand), (4) four inoculated seedlings were assigned as control for the natural abundance of ^13^C_4_-Asn and received basic nutrient solution containing ^14^N-KNO_3_ (3 µmol N kg^-1^ sand) and ^12^C_4_^14^N-Asn (3 µmol N kg^-1^ sand). Seedlings were watered daily and grown for 60 days in the climate chamber.

### Experimental setup

60 DAS clover plants from the testing group (n=96) were randomly assigned into 24 treatments giving four biological replicates per treatment (Fig 1b). Based on current understanding of amino acid uptake and interaction with IN, three general mechanisms were proposed: (1) a single N source: ON or IN independent of the presence of the other one, (2) IN/ON ratio, (3) neither IN or ON, but the total N. To support or reject these modes of control of Asn uptake, we designed an experimental space that systematically covers a matrix of ON (Asn) and IN (KNO_3_) conditions (Figure 1). For one week, plants were treated with 24 nutrient solutions containing different combinations of ^15^N-labeled Asn and KNO_3_ (µmol N kg^-1^ sand). Treatment solutions contained 1) constant Asn concentration with increasing dose of KNO_3_, 2) constant KNO_3_ concentration with increasing Asn dose, 3) Asn and KNO_3_ supplied at ratios of 1/3, 1/2, 2/3, 4) total N corresponding to equimolar concentration of Asn and KNO_3._ Control plants received respective basic nutrient solutions. On day seven, Asn in all the treatment solutions was replaced by the labeled ^13^C_4_-Asn (98 atom% ^13^C) to determine the uptake rate of Asn. Plants were immersed in the solution for 60 min. After that time, plants were taken out and the shoots were cut off. Roots and shoots were thoroughly washed in 0.5 mM CaCl_2_, dried with a paper towel, and frozen in liquid N.

**Fig 1. Experimental design.** (a) Hypothetical models to explain regulation of Asn uptake, model 1 represents a single N source: ON or IN independent of the presence of the other one, model 2 represents ON/IN ratio, and model 3 represents the total N. (b) Matrix of 24 treatments to investigate possible regulation effects on Asn uptake according to model 1 (solid line), 2 (dotted line), and 3 (dashed line), each point on the graph represents one treatment (n = 4), the x-axis indicates the concentration of organic N (Asn), the y-axis indicates the concentration of inorganic N (KNO_3_)

### Analysis

#### ^15^N, total N, and C analysis

Freeze-dried roots and shoots were ground (Geno/Grinder 2000; CertiPrep. Metuchen, NJ 08840) and analyzed for total N and C, and ^15^N-enrichment (EA-IRMS EA, Thermo Fisher Scientific, Bremen, Germany).

#### ^13^C_4_-Asn uptake and amino acid analysis

2 mg of the freeze-dried (72 hr) ground plant material was extracted in a 1-ml extraction mixture of chloroform, methanol and water (1:3:1, v:v:v) in Sarstedts Eppendorf tubes (Sarstedt AG & Co, Nümbrecht, Germany). One metal bead (a 3-mm tungsten carbide bead) was added to each tube. All tubes were shaken (1300 rpm, 3 min) in a 2010 Geno/Grinder (SPEX Sample Prep., Metuchen, NJ 08840). After shaking, the metal beads were removed, and the tubes were centrifuged (10,000 rpm, 4°C, 10 min). From each of the root and shoot extracts, 200 µl was transferred to 0.1 ml inserts (Mikrolab Aarhus A/S, Denmark) placed in Eppendorf tubes. The rest of the supernatant was stored at - 18°C. To each of the extracts, 25 µl of internal standard (norvaline, 0.5 µM) was added and then all evaporated to dryness in a SpeedVac Concentration (Savant, Fisher Scientific, Denmark). The dry extracts were then re-suspended in 20 µl of 20 mM HCl. Extracts were derivatized using an AccQ•Tag Ultra Derivatization Kit (Waters Corp.) according to the manufacturer’s protocol and analyzed using an HPLC (Agilent) coupled to a mass spectrometer (4500 QTRAP (Sciex)) using electrospray ionization in positive ion mode (ion voltage of 4500 eV). Full details of the method can be found in (9). The presence of compounds in the samples was confirmed by comparing the retention times and MRM transitions with reference standard compounds (S1 Table). The compounds were analyzed using Sciex Analyst 1.6.2 software. Calibration curves (0.001–1 pmol μL^-1^) of the unlabeled standards were prepared, and the peak area of each standard was plotted against the standard concentration. A linear function was applied to the calibration curves and used to calculate the concentrations of the amino acids in the samples,

### Calculations and statistics

The proportion of clover N derived from symbiotic N_2_-fixation was calculated using the ^15^N isotope dilution method (21):

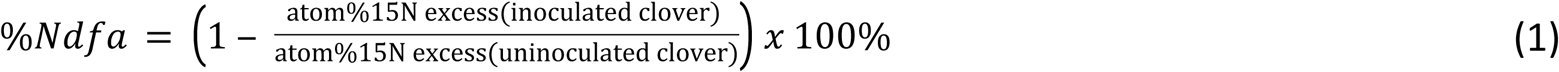

Where the atom% ^15^N excess of inoculated and uninoculated clover were calculated as the atom % ^15^N for the clovers receiving ^15^N-enriched nutrient solution substracted the atom % ^15^N for the respective inoculated or uninoculated clover control receiving un-enriched nutrient solution.

No labeled ^13^C_4_-Asn was detected in the unlabeled clover roots and shoots therefore, no corrections for the natural abundance was made when calculating the excess of labeled amino acid. The net uptake rate of intact ^13^C_4_-Asn (μmol g^-1^ root DW h^-1^) by clover was calculated by adding the amount (μmol) of ^13^C_4_-Asn in the shoots and roots and then dividing by the dry weight of the roots (g). A two-way Anova was conducted to compare the main effects of inorganic and organic N dose and the interaction effect on the Asn uptake rate, amino acid concentration, and clover performance followed by Tukey’s test. The model assumption of normality was tested using the Shapiro-Wilk test, and the assumption of equal variance was tested using Levene’s test and a plot of the residuals against the fitted values. Relationships between ^13^C_4_-Asn uptake rate and Asn concentration in the roots and shoots were tested by Pearson’s correlation analysis. To visualize and characterize the major sources of variability in the samples form the different ON, IN, and total N treatments, Principal Component Analysis (PCA) was applied to autoscaled data of the amino acid concentrations, ^13^C_4_-Asn uptake, total N and C. The data were analyzed using R Studio 3.1.1.

## Results

We did not find evidence that the obtained results were related to the effect of different ratios of N forms (S2 Table) which demonstrates that the IN/ON ratio model is likely not a major regulatory mechanism, at least under the conditions tested. Thus, we have focused the presentation on the effect of IN, ON, and total N.

### General clover performance

Clover was actively fixing N_2_ in all the treatments with the majority of clover N obtained from N_2_-fixation (i.e. %Ndfa ranging from 89-97%) (S3 Table), but no significant changes in biomass, root-shoot ratio, total N and C were found (S4-S7 Tables).

### ^13^C_4_-Asn uptake

The ^13^C_4_-Asn uptake was markedly affected by the different N doses. There was a significant interactive effect of IN and ON dose on the ^13^C_4_-Asn uptake rate (*F*_9,48_ =18.16, *p*<0.05) (Table 1). Specifically, the uptake rate was significantly greater at the lowest IN and ON doses than at increasing IN and ON doses (Fig 2, Table 2). A decreasing pattern of net uptake rate of ^13^C_4_-Asn (*p*<0.05) was observed with increasing total N dose. At low total N (3 IN and 3 ON) the uptake (16.1 nmol g^-1^ root DW) was eight times greater than the uptake (2.1 nmol g^-1^ root DW) at higher total N (30 IN and 30 ON). The interaction between ON dose and IN dose in regulating ^13^C_4_-Asn uptake, was found to respond to IN in an ON-dependent manner. It was observed that at 3 and 30 ON (*p*<0.05) as well as at 20 ON dose (*p*>0.05), the net uptake rate decreased from 50-80% alongside increasing IN dose (Fig 2, Table 2), while the effect of ON on the ^13^C_4_-Asn uptake was not as clear. Namely, when clover plants were exposed to 3 IN dose, the net uptake rate of ^13^C_4_-Asn markedly declined, and then increased together with increasing ON dose, while for the remaining 10, 20 and 30 IN doses this tendency was not shown.

**Table 1.** Two-way Anova for the effects of IN and ON dose (µmol kg^-1^ sand) on Asn uptake rate (nmol g^-1^ root DW), Asn concentration in roots (µmol g^-1^ DW) and Asn concentration in shoots (µmol g^-1^ DW) of white clover.

| Effect | Uptake rate ($\text{nmol g}^{-1}$ root DW) | | | | Asn in roots ($\mu\text{mol g}^{-1}$ DW) | | | Asn in shoots ($\mu\text{mol g}^{-1}$ DW) | | |
| --- | --- | --- | --- | --- | --- | --- | --- | --- | --- | --- |
|  | *df* | SS | *F* | *P* | SS | *F* | *P* | SS | *F* | *P* |
| IN dose | 3 | 310.5 | 43.9 | < 0.001 | 9478 | 31.7 | < 0.001 | 13554 | 35.1 | < 0.001 |
| ON dose | 3 | 192.6 | 27.2 | < 0.001 | 996 | 3.3 | <0.05 | 5331 | 13.8 | < 0.001 |
| IN dose x ON dose | 9 | 385 | 18.1 | < 0.001 | 2562 | 2.8 | <0.01 | 6119 | 5.2 | < 0.001 |
| Error | 48 | 113.1 |  |  | 4778 |  |  | 6171 |  |  |

*df* (defrees of freedecom); SS (sum of squares)

**Table 2.** Uptake rate. Uptake rate of ^13^C_4_-Asn (nmol g^-1^ root DW) by the clover treated with different doses of ON and IN (µmol kg^-1^ sand).

| IN dose | ON dose |  |  |  |
| --- | --- | --- | --- | --- |
|  | 3 | 10 | 20 | 30 |
| 3 | 16.1 ± 1.3 <sup>e</sup> | 2.5 ± 1.6 <sup>ab</sup> | 5.6 ± 3.6 <sup>bcd</sup> | 7.6 ± 2.7 <sup>cde</sup> |
| 10 | 8.9 ± 1.5 <sup>de</sup> | 7.4 ± 1.7 <sup>cde</sup> | 2.2 ± 0.4 <sup>ab</sup> | 1.6 ± 0.7 <sup>a</sup> |
| 20 | 2.6 ± 1.1 <sup>ab</sup> | 2.8 ± 0.5 <sup>abc</sup> | 2.0 ± 1.0 <sup>ab</sup> | 4.4 ± 0.6 <sup>abcd</sup> |
| 30 | 3.0 ± 0.8 <sup>abc</sup> | 2.3 ± 1.0 <sup>ab</sup> | 2.7 ± 0.6 <sup>abc</sup> | 2.1 ± 1.4 <sup>ab</sup> |

Values with different superscripts are significantly different ( $p < 0.05$ ). Data are mean ± sdev

**Fig 2.**
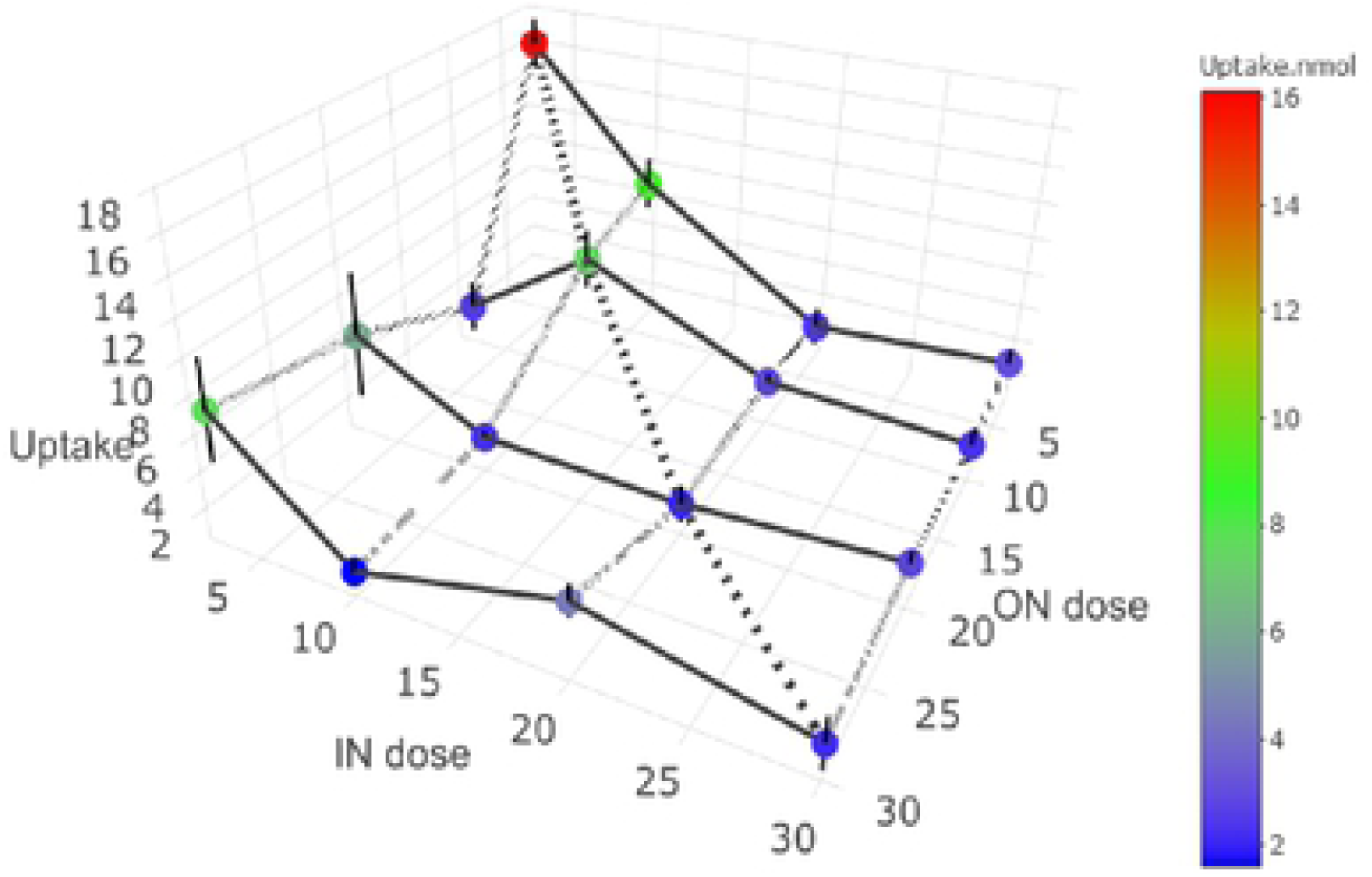
Uptake rate. Uptake rate of ^13^C_4_-Asn (nmol g^-1^ root DW) by the clover treated with different doses of ON and IN (µmol kg^-1^ sand). Data are mean ± sdev

### Asn concentration in the roots and shoots

The Asn concentration in white clover shoots and roots was significantly affected by a combined effect of IN and ON doses (*F*_9,48_ =2.86, *p*<0.05) (Table 1). In the roots, the highest Asn concentrations were found at the lowest IN dose irrespective of the ON dose with 50-60% decreases in root Asn concentration as IN dose increased (Table 3). However, the pattern for Asn concentration was not clear with increasing ON dose, where Asn concentration was observed to variably rise or decline. The interactive effect of ON and IN doses (*F*_9,48_ =5.29, *p*<0.05) was also shown on the Asn concentration in the shoots (Table 1). The highest Asn content was observed at the lowest IN and ON dose, which decreased alongside increasing ON and IN doses resembling the data of the ^13^C_4_-Asn uptake rate (Table 4). At the 3 IN dose, Asn concentration initially decreased, but then increased together with increasing ON dose (*p*<0.05). Interestingly, increasing total N doses significantly (*p*<0.05) reduced both root and shoot Asn concentrations in a similar manner like in case of ^13^C_4_-Asn uptake rate. We found positive correlations between the ^13^C_4_-Asn uptake rate and the Asn concentration in the roots and shoots, respectively, with the strongest correlation found for shoots (R = 0.83) (Fig 3).

**Table 3.** Asn concentration in the clover roots. Asn concentration (µmol g^-1^ DW) in the roots of clover treated with different doses of ON and IN (µmol kg^-1^ sand).

| IN dose | ON dose |  |  |  |
| --- | --- | --- | --- | --- |
|  | 3 | 10 | 20 | 30 |
| 3 | 59.3 ± 19.5<sup>bc</sup> | 41.8 ± 6.5<sup>ab</sup> | 58.4 ± 5.3<sup>bc</sup> | 73.6 ± 5.7<sup>c</sup> |
| 10 | 36.2 ± 9.4<sup>ab</sup> | 37.1 ± 7.7<sup>ab</sup> | 24.7 ± 12.3<sup>a</sup> | 34.7 ± 6.2<sup>ab</sup> |
| 20 | 25.6 ± 6.0<sup>a</sup> | 23.5 ± 11.8<sup>a</sup> | 29.0 ± 2.5<sup>a</sup> | 43.3 ± 9.4<sup>ab</sup> |
| 30 | 29.2 ± 8.2<sup>a</sup> | 32.5 ± 14.8<sup>a</sup> | 26.4 ± 8.9<sup>a</sup> | 23.6 ± 11.6<sup>a</sup> |

Values with different superscripts are significantly different ( $p < 0.05$ ). Data are mean ± sdev

**Table 4.**
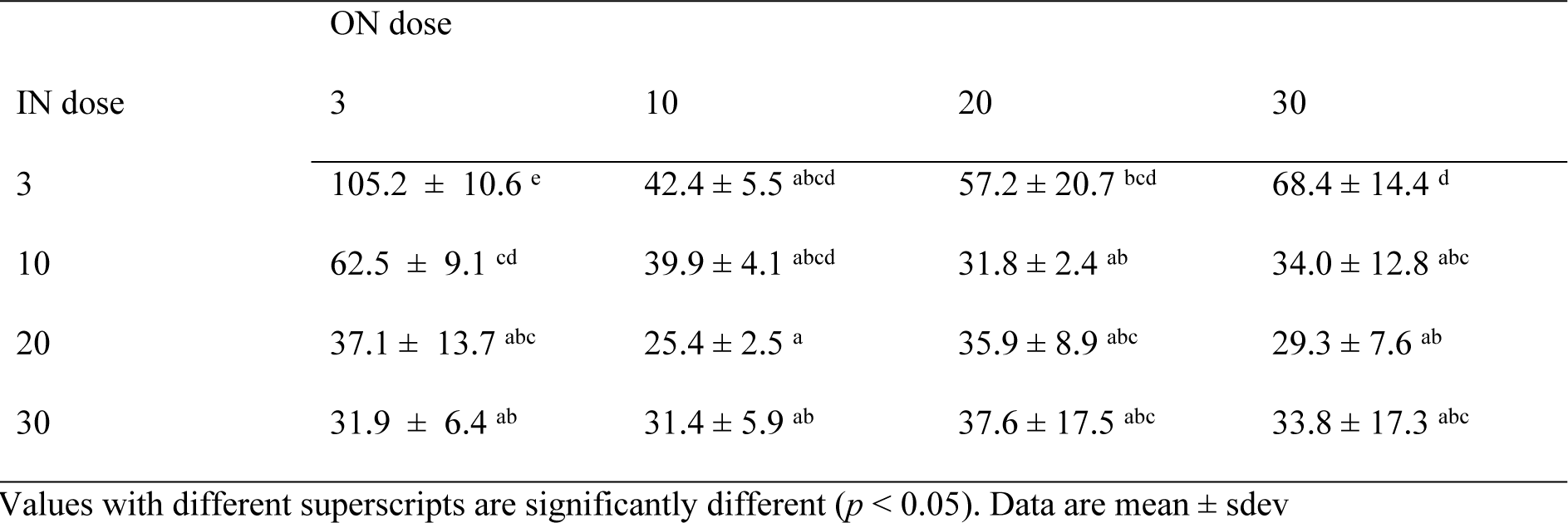
Asn concentration in the clover shoots. Asn concentration (µmol g^-1^ DW) in the shoots of clover treated with different doses of ON and IN (µmol kg^-1^ sand).

**Fig 3.**
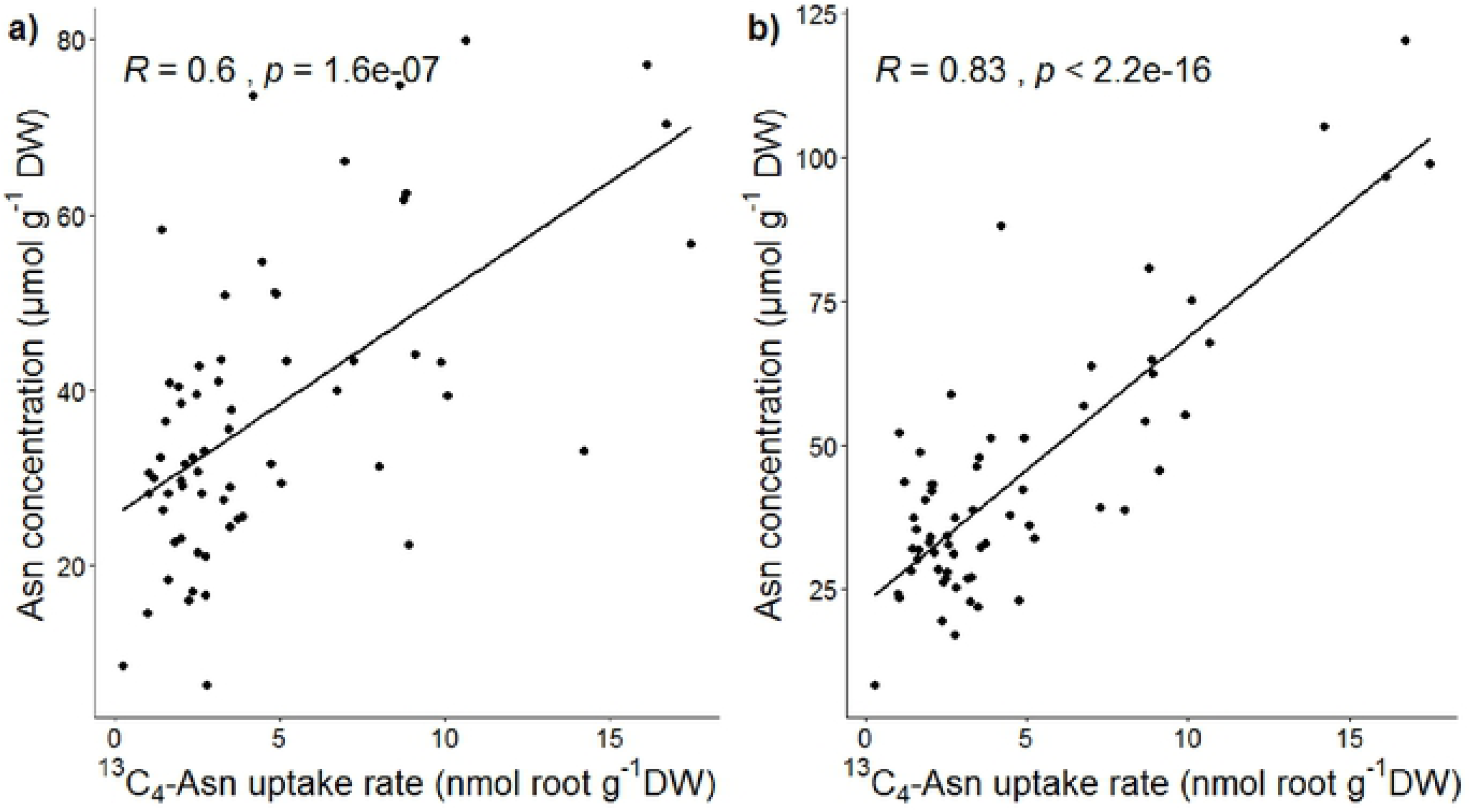
Correlation analysis. Pearson correlation analysis between ^13^C_4_-Asn uptake rate (nmol g^-1^ root DW) and Asn concentration (µmol g^-1^ DW) in the roots (a) and shoots (b).

### Correlations between internal amino acid concentrations

PCA of amino acid concentrations, ^13^C_4_-Asn uptake rate, total N and C revealed different groupings depending how objects were assembled in sets. No groupings was found when each singular treatment (one ON and one IN dose) was marked as a separate set (S1 Fig). On the contrary, two groupings related IN and total N dose were revealed. In the shoots, the groups related to the lowest IN (Fig 4a) and Total N (Fig S2b) doses were characterized by higher concentrations of most the non-essential amino acids: Asn, Asp, Glu, Gln, Cys, Pro, Gly and Ala, as well as ^13^C_4_-Asn uptake rate, whereas the groups related to the higher IN and total N doses contained more of the essential amino acids: Thr, Val, Ile, Leu, Phe, Tyr, Trp and Met. The same pattern was shown in the roots (S2a Fig, S3 Fig). No separation related to the ON was observed (Fig 4b).

**Fig 4.**
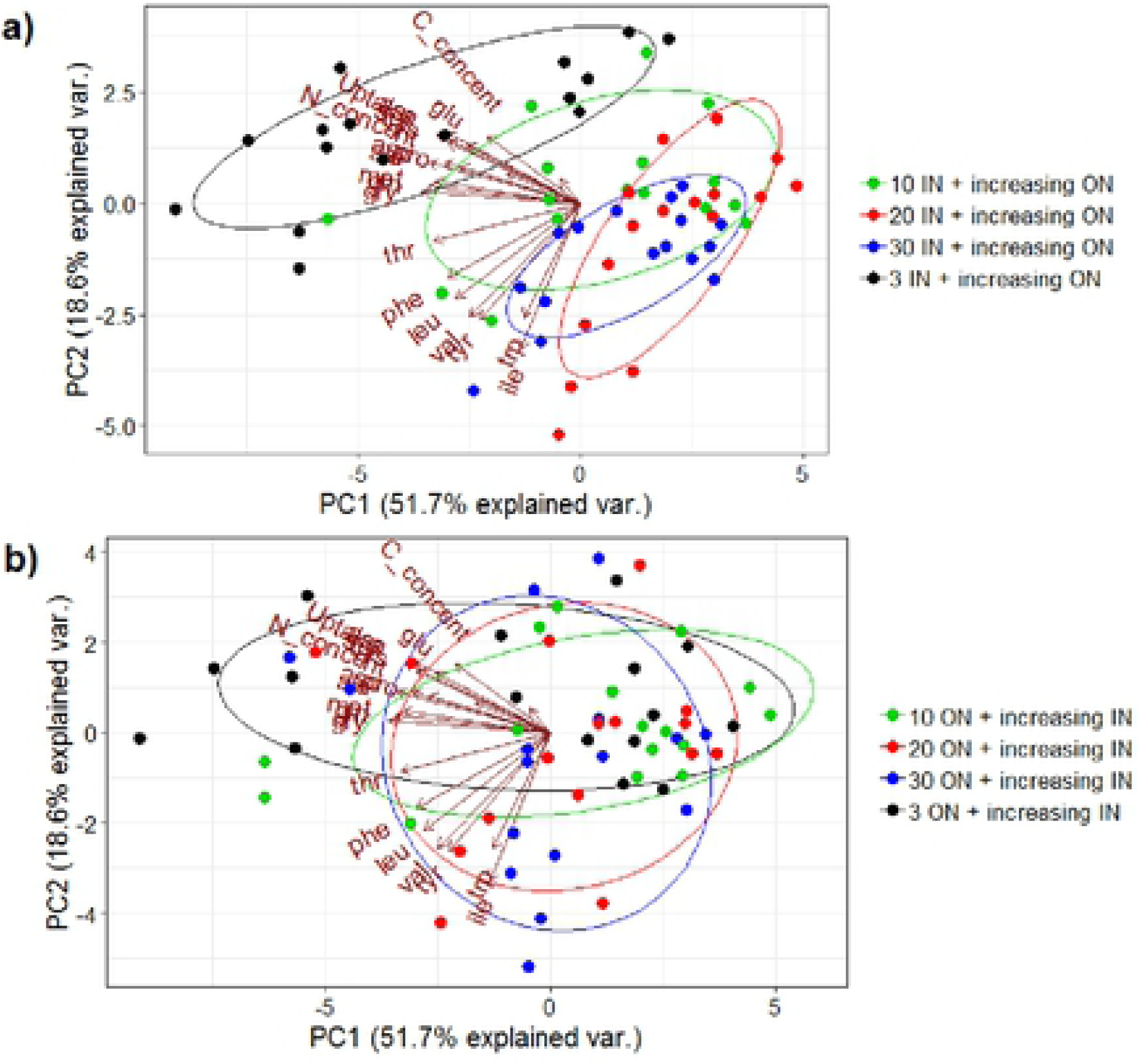
PCA analysis. PCA analysis of amino acids, ^13^C_4_-Asn net uptake rate, total N and C content in clover shoots treated with (a) IN and increasing doses of ON (µmol kg^-1^ sand), (b) ON and increasing doses of IN (µmol kg^-1^ sand).

## Discussion

### ^13^C_4_-Asn uptake is restricted by external IN, but not ON

We found that one week of exposure to increasing, yet low, IN and ON concentrations reduced the ^13^C_4_-Asn uptake rate in white clover by a factor of seven; with the ^13^C_4_-Asn uptake rates being about an order of magnitude lower than previously reported for white clover grown in hydroponics (3) and under soil conditions (9). N uptake is affected both by the internal N status of the plant and the external co-occurrence of N sources (22). In clover no changes in the total N were observed, when exposed to different doses of ON, IN, and total N. Thus, changes in the ^13^C_4_-Asn uptake rate were more related to the presence of external N, with decreasing uptake along the increasing total N and increasing IN at the low ON dose (Table 2). Decreased uptake of amino acid in the presence of IN was also found by (22), where the uptake of arginine by scots pine (*Pinius sylvestris* L.) was twice as high when provided alone compared to when supplied with NO_3_^-^. The reduced uptake of ^13^C_4_-Asn with increasing IN doses demonstrates that assimilation of amino acids is less relevant to the clover under those conditions. However, a greater uptake of the amino acid under limiting IN availability could also point to that clover have a high flexibility to fulfill optimal N nutrition under various conditions. Such properties were for example reported for tree species (15), tundra plants (23) or deciduous and coniferous taiga forest (24) growing in natural ecosystems where ON nutrition is important due to slow mineralization rate. Increasing ON doses resulted in a more complex effect on the ^13^C_4_-Asn uptake (Table 2). Some studies documented a downregulation of amino acid uptake by IN (12), whereas other reported an amino acid uptake to be concentration-dependent and that increasing amino acid concentration results in increased uptake rates. This was observed for wheat (*Hordeum vulgare* L.) supplied with glycine (2-30 µ*M*) (25), and spruce (*Fagus sylvatica*) supplied with glutamine (1 µM-10 m*M*) (26). The reports of increasing amino acid uptake rates with increasing amino acid concentrations all come from studies of non-legumes, whereas the present results indicate that increasing amino acid concentrations has the opposite effect on amino acid uptake in legumes. We conclude that it was a complex interaction due to the co-occurrence of different N forms, rather than the presence of one N form that affected clover amino acid uptake. Similar to the effect of increasing IN doses a reduced ^13^C_4_-Asn uptake rate was found for increasing total N doses (equimolar concentration of IN and ON) (Table 2), which point to a greater influence of IN than ON in the regulation of amino acid uptake in clover.

### ^13^C_4_-Asn uptake correlates with the internal Asn concentration

Similar to ^13^C_4_-Asn uptake rates, the Asn concentration in white clover shoots and roots was significantly influenced by exposure to increasing IN, ON, and total N doses; with the Asn concentration declining along with increasing external total N (Tables 3 and 4). White clover in this study would have three sources of N available: 1) N from NO_3_^-^ uptake, 2) N from Asn uptake, and 3) N from N_2_-fixation. It is clear that with lower ^13^C_4_-Asn uptake rates at increasing external N doses either NO_3_^-^ uptake or N_2_-fixation must have been upregulated to maintain plant N status. However, %Ndfa was stable and unaffected by the different N doses (S3 Table). Therefore, it is more likely that NO_3_^-^ uptake was increased with increasing external IN and total N doses as the present N_2_-fixation paradigm states that increasing external NO_3_^-^ availability reduces the N_2_-fixation activity (27). For instance, Sulieman, Schulze (28) grew *Medicago truncatula* under high NO_3_^-^ (5 m*M*) and observed a significant reduction in nitrogenase activity along with Asn accumulation in nodules. The decrease in enzymatic activity was further associated with Asn build-up participating in a negative N-feedback and inhibiting nitrogenase in response to excessive NO_3_^-^. In the present study the decreasing Asn concentration in shoots with increasing IN and total N doses could therefore be explained by Asn loading into nodules to reduce N_2_-fixation (Table 4). However, we did not see a greater Asn accumulation in the roots, which were sampled together with nodules, nor did we observe a decrease in the N_2_-fixation. This result is unusual because a negative feedback system is common for most of the legume plants (29). In retrospect, it would have been very useful if we had sampled nodules separately to measure nodule Asn concentration, because the present findings on the relation between external N doses and internal Asn concentrations does not seem to be directly in line with the hypothesis of Asn being the key in internal regulation of N_2_-fixation in legumes. Alternatively, other pathways in which Asn is utilized for synthesis of other amino acids could be a reason for a declining Asn content (18, 30), as we observed an increasing tendency in the concentration of Phe, Thr, and Tyr alongside increasing total N (S4 Fig). However, more research would be needed to establish this link. We furthermore were puzzled by finding a positive correlation between shoot Asn concentration and the uptake rate of ^13^C_4_-Asn across the external IN and ON doses, which points to that the two are connected. Although we cannot here deduce whether uptake rate controls internal concentration or vice versa.

### Internal amino acid composition is affected by external N doses

In parallel to ^13^C_4_-Asn uptake rate and internal Asn concentration, external IN and total N doses changed the concentrations of amino acids in the roots and shoots. At low N, we found a dominance of non-essential amino acids including Asn, whereas increasing external N changed the amino acid profile towards essential amino acids (Fig 4). At the same time, the total N content remained unchanged (S7 Table). This is in line with observations made by Ferreira, Novais (18), who concluded that free-amino acids show a greater promise than total N in understanding the effect of external N on the plant N status, in that amino acid content can respectively increase or decrease in stress conditions without any changes in the total N. Although no studies could be found on the effect of external N on amino acid profile in legumes, (31) reported a decrease in the content of non-essential amino acids in response to high N for maize (Zea mays L.). They hypothesized that decline of the non-essential amino acids was due to deficiency of carbon skeletons for the assimilation of NH_4_^+^. Perhaps a decreased content of Asn in our study (Tables 3 and 4) could also be linked to its metabolization, so that Asn carbon skeletons could be precursors for the synthesis of other amino acids. In that context our findings support that not only the amount of amino acids, but also information on the composition of amino acids is needed to determine to what extent the plant is N stressed and how the plant signals N demand between root and shoot. Thus, our findings demonstrate that the external organic and inorganic N affects the accumulation of certain amino acids in clover, which could help in further understanding how the plant senses various N stress conditions and circulates amino acids between roots and shoots (18, 32). Soil inorganic and organic N status could therefore be used as an indicator for nutritional quality of protein content in forage legumes. Furthermore, our results on shoot and root amino acid composition could also be relevant in understanding how legumes regulate N_2_-fixation activity as e.g. Glu, Gln, and Pro (33-35) have been found related to the N_2_-fixation regulation in addition to Asn. Indeed, we found that Glu, Gln, and Pro were all related to both Asn and ^13^C_4_-Asn uptake rate at the low external N doses (Fig 4), which would be the conditions where we would expect the greatest N_2_-fixation activity in the white clover. Thus, in future studies on the impact of external N concentrations and forms it would be relevant to investigate how increasing soil N affects the amino acid profile and N_2_-fixation activity in legumes and N_2_-fixation activity in legumes.

### Conclusion

In conclusion, a complex interaction due to the co-occurrence of different N forms, rather than the presence of one N form affected clover amino acid uptake; with increasing external IN and total N concentrations reducing ^13^C-Asn uptake rates. In addition, increasing total external N concentrations affected both Asn and amino acid profiles indicating that plant amino acid profiles may be a good indicator for plant N status. Interestingly, there was a positive correlation between ^13^C-Asn uptake rate and shoot Asn concentration, although further studies are needed to elucidate whether this link is controlled by the external N concentrations or internal Asn content.

## Acknowledgements

The study was financially supported by The Independent Research Fund Denmark – Technology and Production (Project no. 1335-00760B). The authors would like to thank Professor Wanda Malek (Maria Curie Sklodowska University in Lublin, Poland) for providing bacterial strain *Rhizobium leguminosarum* bv. trifolii TA1 for the plant inoculation.

## Supporting information

**S1 Table**. Mass-to-charge ratio (m/z) used in selected ion monitoring and retention times

**S2 Table** Uptake rate of ^13^C_4_-Asn (nmol g^-1^ root DW) and Asn concentration (µmol g^-1^ DW) in the roots and shoots in the clover treated with different doses of ON and IN (µmol kg^-1^ sand) supplied at different ratios. Data are mean ± sdev

**S3 Table** %Ndfa in clover treated with different doses of ON and IN (µmol kg^-1^ sand). Data are means ± sd

**S4 Table** Total biomass (g) of clover treated with different doses of ON and IN (µmol kg^-1^ sand). Data are means ± sd

**S5 Table** Root:shoot ratio of clover treated with different doses of ON and IN (µmol kg^-1^ sand). Data are means ± sd

**S6 Table** Total C (g g^-1^ DW) of clover treated with different doses of ON and IN (µmol kg^-1^ sand). Data are means ± sd

**S7 Table** Total N (g g^-1^ DW) of clover treated with different doses of ON and IN (µmol kg^-1^ sand). Data are means ± sd

**S1 Fig.** PCA of amino acids, ^13^C_4_-Asn net uptake rate, total C and N content in clover shoots (a) and roots (b) treated with different doses of ON and IN.

**S2 Fig.** PCA analysis of amino acids, ^13^C_4_-Asn net uptake rate, total N and C content in clover roots (a) and shoots (b) treated with increasing total N dose.

**S3 Fig.** PCA analysis of amino acids, ^13^C_4_-Asn net uptake rate, total N and C content in clover roots treated with (a) IN and increasing doses of ON, (b) ON and increasing doses of IN.

**S4 Fig.** Phenylalanine, tryptophan, and tyrosine concentration (µmol g^-1^ DW) in the shoots (a) and roots (b) at total N (equilimolar concentration of ON and IN µmol kg^-1^ sand DW).

